# Extensive Dysregulation of SLK Splicing in Cancers Impacts Metastasis

**DOI:** 10.1101/2022.10.28.514146

**Authors:** Ying-Qun Yang, Yue Hu, Si-Rui Zhang, Jie-Fu Li, Jia-Wen Guan, Wen-Jing Zhang, Yu Sun, Xiao-Yan Feng, Jing Sun, Yun Yang, Zefeng Wang, Huan-Huan Wei

## Abstract

RNA splicing control is a pivotal aspect of gene regulation and is closely associated with cancer development. From a pan-cancer transcriptome investigation in the splicing layer, we discovered a critical cancer-associated alternative splicing (AS) event at exon 13 of SLK which produces two isoforms, SLK-L and SLK-S. The splicing is dramatically shifted towards SLK-L across multiple prevalent cancer types. We demonstrated that SLK-L plays an essential role in cancer development, especially in metastasis both in cells and in mice, whereas splicing toward SLK-S inhibits cancer development. RNA-seq revealed the two SLK isoforms play different roles in pathways related to cell migration. Furthermore, different SLK isoforms demonstrate varying binding affinities to certain cell junction markers, in part indicating the AS of SLK contributes to cancer cell migration. In addition, the splicing factor Rbfox2 was identified to specifically inhibit the inclusion of exon 13 by binding intron 12 of SLK. Collectively, our study innovatively uncovers the biological consequences and underlying mechanisms for one of the most mis-spliced genes in cancer, highlighting its potential significance in cancer diagnosis and treatment.

## Introduction

RNA splicing control is a critical layer of gene regulation in addition to transcriptional and epigenetic regulation. Extensive alteration of splicing is considered an established hallmark of cancers ^1–8^. Studying splicing dysregulation in cancers is a challenge due to the large noise, as well as the diverse and complicated mechanisms. We previously established a transcriptome-wide identification of cancer-specific splicing events across several tumors through analysis of transcriptome sequencing data from The Cancer Genome Atlas (TCGA) ^9^. This method was utilized to analyze multiple prevalent forms of cancer that have an adequate number of paired normal samples in this study. As a result, we identified a critical cancer-associated splicing event at the exon 13 of STE20-like protein kinase (SLK), which produces two isoforms, SLK-L and SLK-S. Impressively, the splicing is dramatically shifted towards SLK-L among thousands of cancer patients across multiple cancer types.

SLK is a serine/threonine kinase and belongs to the five germinal center kinase families. It contains several conserved domains, including a kinase domain, a predicted central unstructured region, and a C-terminal coiled-coil domain. Previous studies discovered that SLK was involved in several cellular processes such as cell cycle progression by phosphorylating PLK1 and promoting G2/M phase transformation ^10, 11^, or activating the phosphorylation of p38 mitogen-activated protein kinase (MAPK) through ASK1 ^12, 13^. Over the twenty years, SLK has emerged as a regulator of the cytoskeleton ^14^. It has been shown to activate the Rho family of GTPases, which in turn regulate cytoskeletal dynamics and cell migration ^15^. It impacts the migration of mouse fibroblasts by directly interacting with LIM domain-binding protein 1 (Ldb1) ^16^, and is involved in the epithelial to mesenchymal transition (EMT) process ^17–19^ and the regulation of focal adhesion turnover ^20^ through its kinase activity ^15^.

Interestingly, several recent studies revealed that the ability of SLK for regulating cellular functions appears to be kinase activity independent, suggesting that it may be an important scaffold for signal transduction pathways^14^. For instance, in fibroblasts, either the expression of an SLK variant lacking kinase activity or depletion of SLK can result in growth arrest, indicating that both kinase-dependent and-independent functions of SLK are necessary ^17^. Functions of Slik, the homolog of SLK in *Drosophila*, are dependent on its proper subcellular localization, which is mediated through its C-terminal ATH domain independent of its activation status ^21^. These results suggested that the alternative 13^th^ exon in the ATH domain of SLK may have important functions. However, the underlying mechanisms remain unknown.

Accumulating evidence has shown that *SLK* plays important role in cellular functions associated with cancer development ^14, 22, 23^. However, few studies report and investigate the splicing switch of *SLK* in cancers. The prevalent AS alteration of *SLK* indicates that the two isoforms may play different roles in cancers. Furthermore, the extensively dysregulated splicing events across cancer types suggest the possibility of universal functional and mechanistic implications. Therefore, uncovering the functions and mechanisms of abnormal splicing of *SLK* in cancers will provide insights into the cancer-associated key AS events with therapeutic potential and clinical values.

## Results

### *SLK* splicing is significantly and extensively altered across multiple common types of cancer

Using the previously established method ^9^, we performed a pan-cancer investigation of global AS dysregulation by comparing the Percent Spliced In (PSI) value between tumor samples and their adjacent normal controls (Fig. 1a). For a reliable comparison, we selected the most prevalent forms of cancer that have a sufficient number of paired normal samples. As a result, we identified a substantial number of splicing events that were differentially expressed in cancer. Our findings indicate that approximately 70% of splicing events are altered across different cancer types (Fig. S1a). This observation further suggests that the shared spliced genes might potentially serve as an underlying biological mechanism that promotes cancer development and progression. Additionally, we observed that such shared genes altering in more than 10 cancer types are enriched in cancer-related migration processes such as actin filament organization, cell junction assembly, and cell-substrate adhesion (Fig. 1b). Of particular interest, we have discovered notable and comprehensive changes in the splicing of SLK gene in 12 different types of cancer for the first time (Fig. 1b and S1a). Besides, in nine cancer types, SLK showed a significant shift toward the long isoform with PSI significantly increased compared to that in normal samples, which indicates AS of SLK play key roles during cancer development (Fig. 1c).

**Figure 1.**
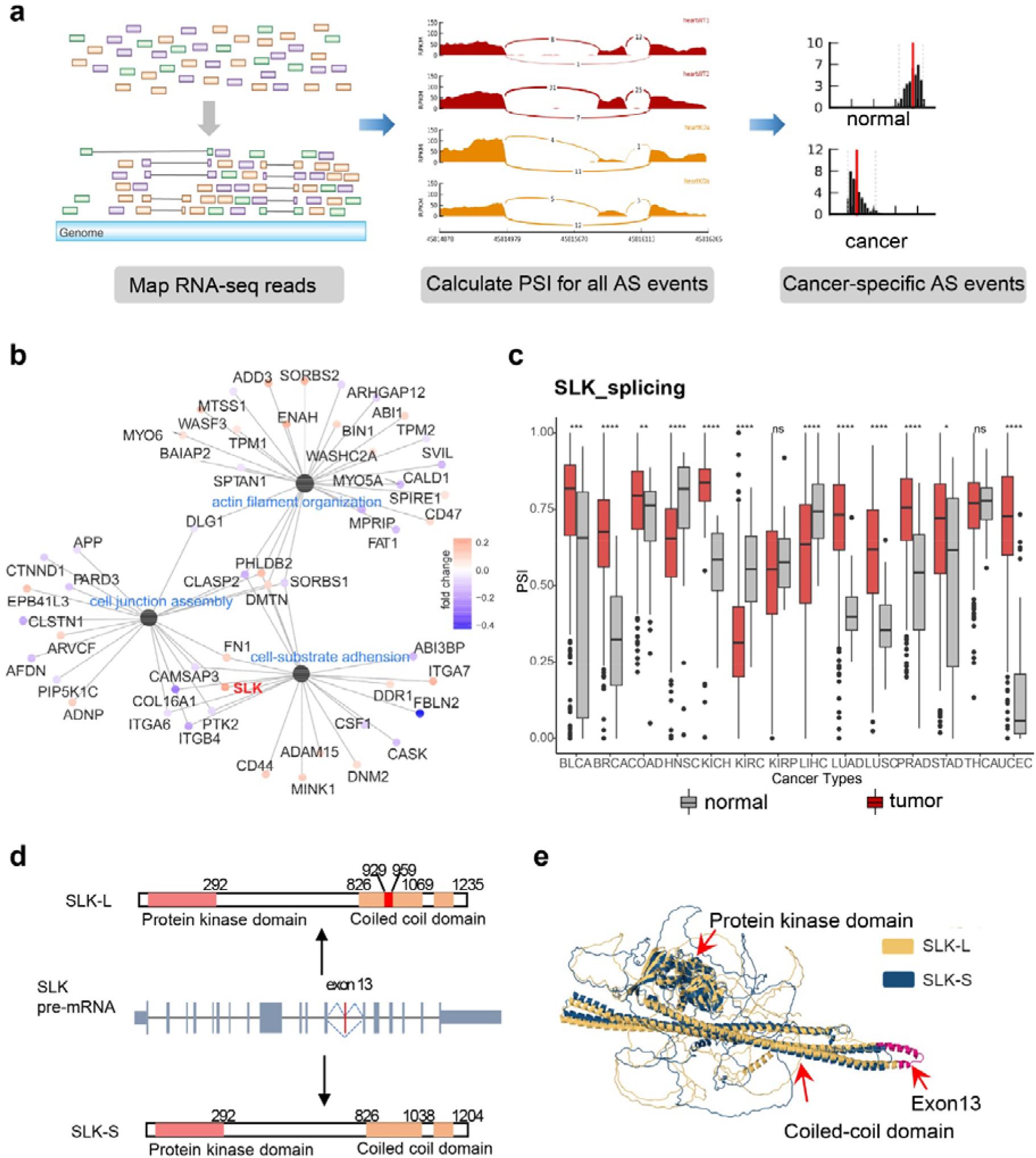
Pan-cancer transcriptome analysis in the splicing layer revealed SLK AS alters dramatically in multiple types of human cancer. **a,** Analysis pipeline to identify AS events differentially spliced in cancers versus normal tissues. **b,** GO enriched network of overlapped AS genes in more than 10 cancer types. **c,** The SLK splicing ratios across various cancer types in the TCGA database. PSI: percent spliced in. **d,** Schematics of human SLK pre-mRNA and protein isoforms produced from SLK AS. the 13^th^ exon is included in SLK-L while skipped in SLK-S. **e,** The comparison of the predicted structure of SLK-L (yellow) and SLK-S (blue) isoform by Alphafold. The alternative spliced exon 13^th^ is highlighted in red.

The human *SLK* gene is located on chromosome 10 and consists of 19 exons. The SLK protein contains the N-terminal kinase catalytic domain (encoded by exons 1 to 8), an interdomain linker, and the C-terminal coiled-coil domain responsible for protein-protein interaction (encoded by exons 11 to 19). The alternative inclusion of exon 13 in *SLK* pre-mRNA produces two isoforms, the long or short isoform (SLK-L and SLK-S), which differs at a fragment of 31 amino acids in the C-terminal coiled-coil domain (Fig. 1d). The comparison of the protein structures predicted by Alphafold (https://alphafold.ebi.ac.uk/, ^24^) demonstrates that the exon 13 encoded sequence locates in the terminus of the long α-helix inside the coiled-coil domain (Fig. 1e), suggesting a kinase-independent activity between SLK-L and SLK-S. The structure of SLK also indicates its potential role as scaffolding protein.

SLK AS event is conserved in mammals (Fig. S1b), indicating functional importance. We further investigated the expression of SLK isoforms in various tissues using Genotype-Tissue Expression (GTEx) data ^25^. The exon count analysis showed a preference for SLK-S expression in tissues with slow self-renewal (e.g., brain), whereas the SLK-L is preferably expressed in tissues with rapid proliferation and self-renewal (e.g., skin). In particular, the non-dividing muscular tissue of the esophagus mainly contains the SLK-S isoform, whereas the fast-renewal mucosa membrane of the esophagus contains the predominant SLK-L isoform (Fig. S1c). Such pattern is largely consistent with the *SLK* splicing changes in tumors that contain many actively dividing cells (Fig. 1c), implying that SLK-L may play a role in promoting tumor development. Moreover, the mechanism of this dysregulation may play general roles across cancer types.

### Alternative Splicing of SLK affects cancer cell proliferation and migration

To further study the function of AS in SLK, we used siRNAs that were designed to specifically target SLK-L (binding to the 13^th^ exon of SLK) or SLK-S (binding to the junction of the 12^th^ and 14^th^ exons) in colon cancer HCT116 cells (Fig. S2a-S2b). The highly invasive HCT116 cells have a normal diploid karyotype with the PSI of the SLK between 0.4-0.7, which is ideal to examine the change of AS ratio in both directions. We found that the cells with SLK-S knockdown showed similar cellular phenotypes compared to the control cells, whereas cells with SLK-L knockdown showed a significant reduction in cell proliferation, anchorage-dependent colony formation, migration, and invasion (Fig. S2c-S2f), suggesting that the two isoforms have distinct cellular functions.

Since siRNA-mediated gene knockdown is transient and often leads to off-target effects, resulting in residual expression of the targeted gene (Fig. s2b). we used CRISPR technology to establish HCT116 stable cell lines that specifically express SLK-L or SLK-S to ensure the reliability and stability of the functional experiments (Fig. 2a-2b). Consistent with the siRNA knockdown results, HCT116 cells with only SLK-S showed a significant reduction in cell proliferation (Fig. 2c), colony formation (Fig. 2d), cell migration (Fig. 2e), and cell invasion (Fig. 2f), again suggesting that SLK-L deficiency may suppress cancer progression. As a comparison, the cells with SLK-S depletion exhibited normal or slightly increased cell growth and cell migration (Fig. 2e-2f, Fig. S2e-S2f), suggesting that SLK-L isoform is required for the robust proliferation and migration of cancer cells.

**Figure 2.**
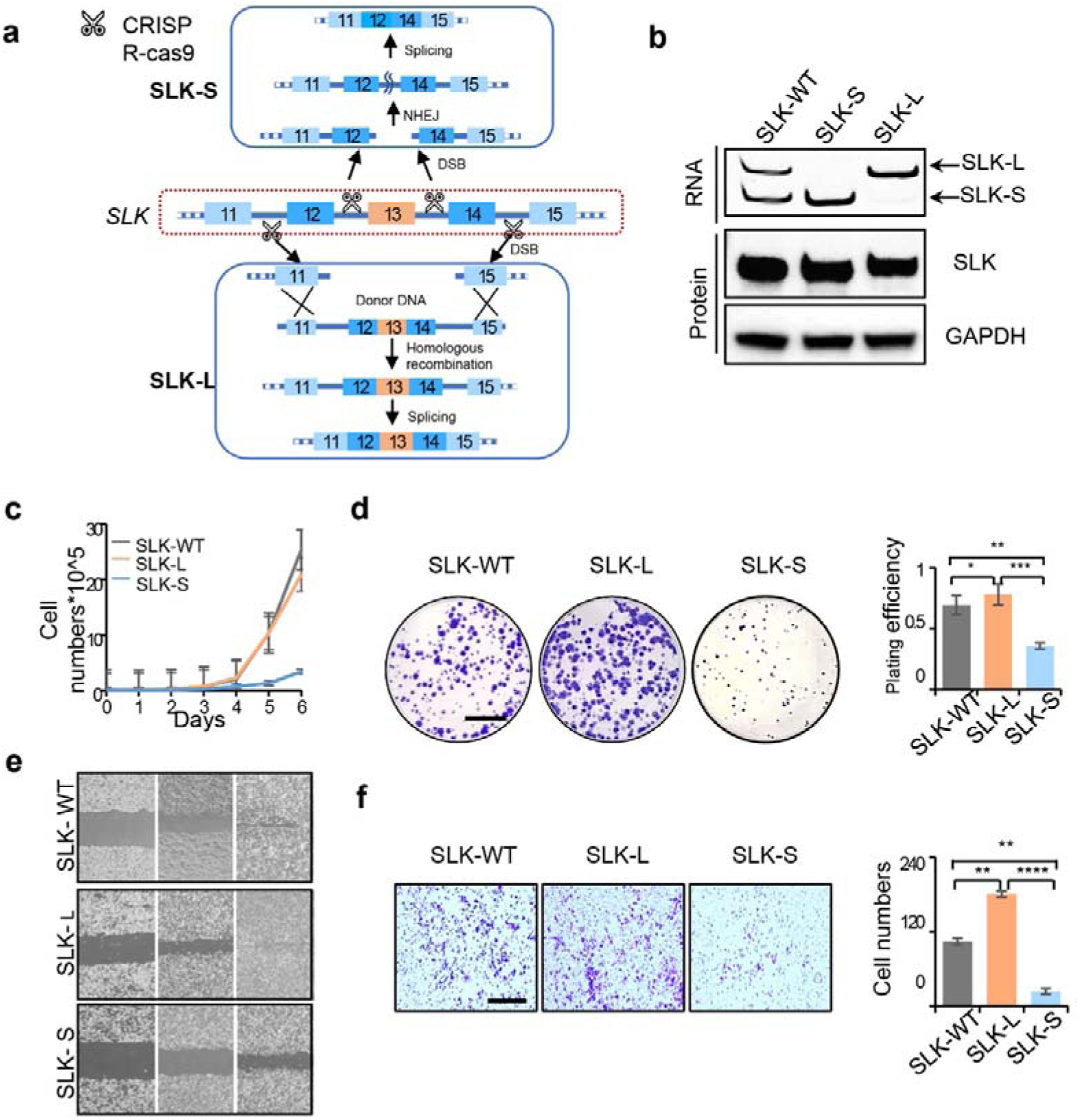
The cancer cells stably express SLK isoforms were generated by CRISPR-cas and exhibited different proliferation and migration abilities. **a,** HCT116 cells expressing different SLK isoforms were constructed by CRISPR. **b,** RNA and protein levels of SLK under CRISPR genome editing. **c-f,** Cancer cell progression was measured from triplicate experiments, with p values calculated by one-way ANOVA and Tukey’s test. **c,** various SLK cells were grown for 6 days, with cell numbers counted every day (n=3) to measure cell growth **d,** anchorage-dependent colony formation of different SLK cell lines (n=3; scale bar:1 cm). The planting efficiency was quantified with ImageJ. **e,** *In vitro* migration of different SLK cell lines was analyzed by wound healing assay (n=3). **f,** cell invasion was measured by transwell assay (n=3; scale bar: 200 µm). Cell number was quantified with ImageJ. *p<0.05, **p<0.01, ***p<0.001, ****p<0.0001, ns, not significant. The chart represents mean±SD.

For the purpose of determine whether the splicing switch of SLK were authentically responsible for the observed cellular phenotypes, we re-expressed the isoforms separately or in combination in the SLK knockout (SLK-KO) HCT116 cells generated with CRISPR-Cas technology (Fig. 3a-3b). Our results showed that cells with re-expressed SLK-L isoform exhibited restored cell proliferation and migration/invasion, while cells with re-expressed SLK-S isoform were similar to SLK-KO cells (see Fig. 3c-3f). These findings provide evidence that the SLK-L isoform promotes the proliferation and migration of cancer cells like an oncogene.

**Figure 3.**
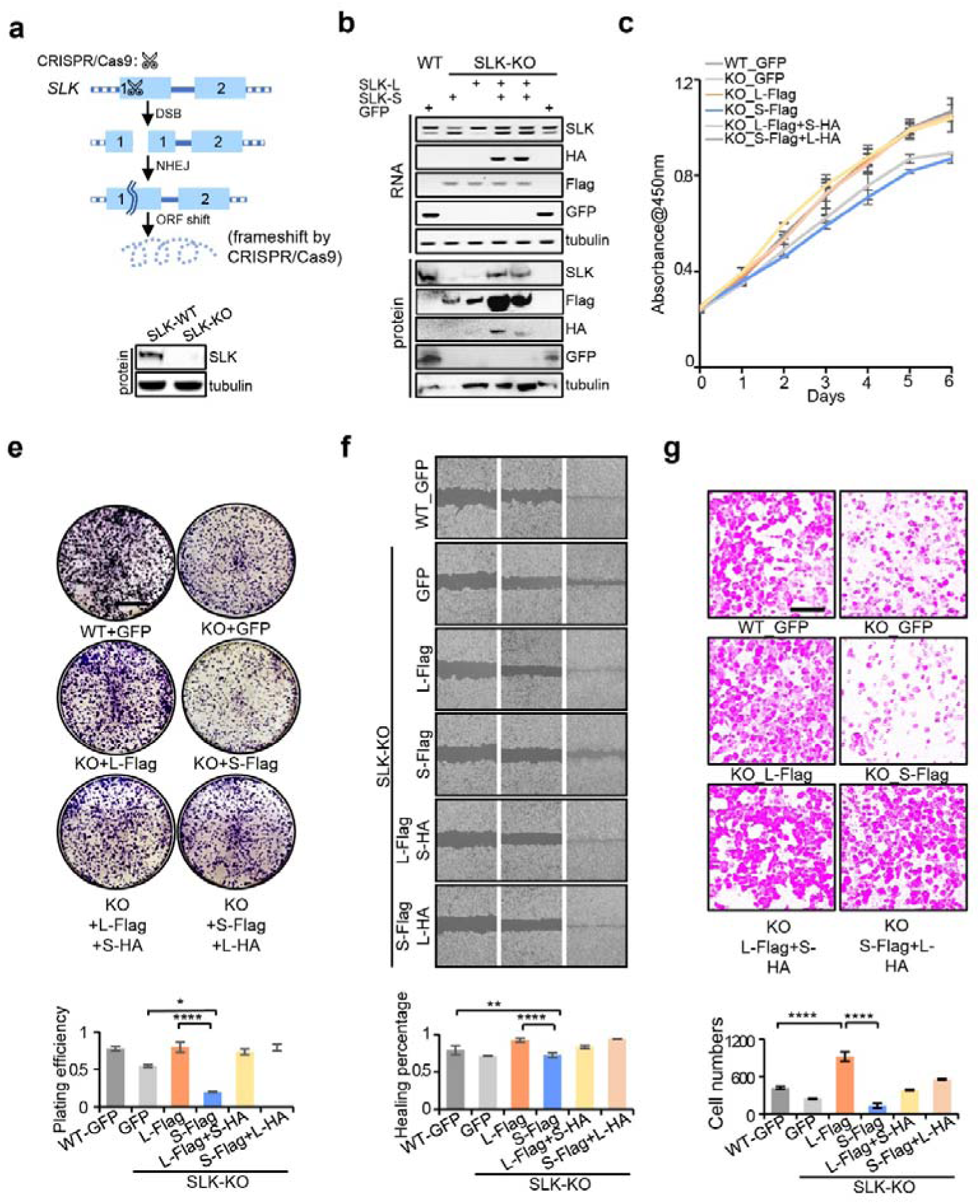
Cancer progression was restored by recovering the expression of SLK isoforms. **a,** Construction of SLK knockout cell line by CRISPR. **b,** SLK isoforms with various tags were re-expressed in SLK knockout cells, with the RNA and protein levels of SLK detected accordingly. **c-g,** Cancer cell development was measured as described in Fig. 2.

### SLK splicing switch affects cancer growth and metastasis in mice

Based on the cell function experiment, we investigated the effect of dysregulated SLK AS on cancer progress *in vivo*. We generated xenograft mice using HCT116 cells expressing SLK-WT, SLK-L, SLK-S, or SLK-KO. Nude mice were inoculated with a low (1×10^6^ cells, Fig. 4a) or high (5×10^6^ cells, Fig. S3A) number of cells in two separate experiments. As expected, the size and weight of the subcutaneous tumors increased rapidly in the SLK-WT and SLK-L groups while they were in smaller size and weight in the SLK-S group (Fig. 4a-4b and Fig. S3a-3c). Moreover, in the high-dosage experiment, the mice inoculated with SLK-WT and SLK-L exhibited lower survival in comparison to the SLK-S group (Fig. S3a). These results indicated that SLK-L contributes to cancer development, and suggested the splicing switch to SLK-S had the potential for cancer suppression. Surprisingly, the SLK knockout cells grew faster than SLK-S cells (Fig. S3a), indicating an undiscovered complementary mechanism upon SLK knockout.

**Figure 4.**
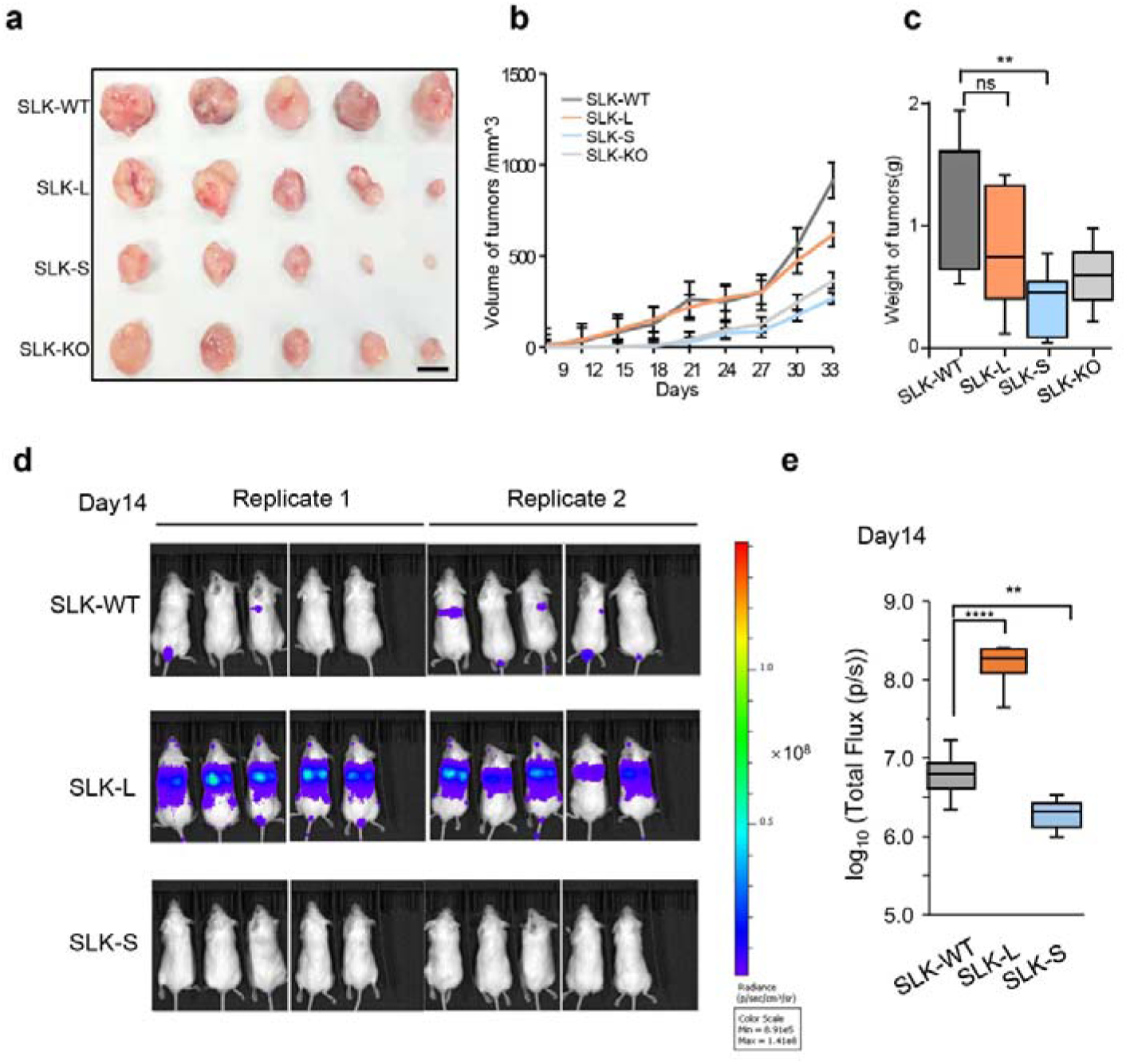
Different groups of SLK cells exhibited distinct impacts on cancer development. **a-c,** Xenograft tumors were generated in BALB/c nude mice by subcutaneous injection with SLK-WT, SLK-L, or SLK-S in **b** and **c**, respectively, with *p* values calculated by Dunnett’s test. **d-e,** Metastasis of SLK cell groups was investigated by the tail vein metastasis model. Different SLK cell groups stably expressing firefly luciferase were injected into the lateral tail vein of immune deficiency mice and the metastasis was monitored by bioluminescence imaging every week post injection **(see also Fig. S3). d,** The bioluminescence images were demonstrated in the two replicates of C-NKG mice on day 14. **e,** Box plot of the mean log_10_ transformed luciferase signal measured for each mouse in d. Statistical significance was calculated using a Student’s *t* test; *p<0.05, **p<0.01, ***p<0.001, ****p<0.0001, ns, not significant.

Interestingly, in situ tumorigenicity assay exhibited that SLK-L cells demonstrated a similar phenotype compared with SLK-WT cells, which is consistent with *in vitro* experiments. However, we noticed that in the *in vitro* experiments, SLK-L cells demonstrated higher metastasis capabilities than SLK-WT cells (Fig. 2 and 3). Considering metastasis is the main cause of cancer-related deaths ^26^, and the emerged functions of SLK as a central regulator of cytoskeletal dynamics ^14, 27^ and cell migration ^28, 29^, we hypothesized that SLK splicing switch has critical effects on cancer metastasis. To testify our hypothesis, we developed a tail vein injection mouse model to examine metastatic colonization and subsequent growth of different SLK cells. Different groups of SLK cells were stably transduced with viral vectors encoding firefly luciferase and injected into mice via the lateral tail vein. The bioluminescence of the tumor cells was monitored and measured to quantify changes in metastatic burden over time. For reliability, the experiment was performed in two immune deficiency mouse strains, the BALB/c nude mice (Fig. S3d) and C-NKG mice (Fig. 4d and S3e). In BALB/c nude mice (Fig. S3d), four in five mice injected with the SLK-L cells demonstrated lung and other organ metastasis before the endpoint (7 weeks post-injection) while only one and zero mice exhibited metastasis in SLK-WT and SLK-S group, respectively (Fig. S3d). In C-NKG mice, lung and other organ metastasis was observed in all SLK groups, with SLK-L cells demonstrating significant higher metastatic burden and fatality than SLK-WT and SLK-S cells (Fig. 4d and Fig. S3e). The faster metastasis formation and growth in C-NKG mice than in BALB/c mice mainly contributed to the higher level of immune deficiency of C-NKG mice, considering that depletion of NK cells in mice increases metastasis ^30^.

### RNA-seq revealed SLK splicing switch influence key components in cell junction and cell migration

Cell and mouse experiments indicated that the different isoforms of SLK had significant impact on cell migration and metastasis. We next sought to determine how splicing switch of SLK isoforms decide cancer progression. Though accumulating studies unveiled that SLK plays functions in EMT and cytoskeleton thereby influences cell migration, there is still a lack of systematic investigation on the functions of SLK and its isoforms. To understand the underlying mechanism of SLK splicing switch in a systematic manner, we further examine the biological functions of SLK isoforms by analyzing RNA-seq data of different SLK cell lines (Fig. 6a). We discovered many differentially expressed genes are enriched in cancer-related biological process, such as the Wnt signaling pathway and tissue migration (Fig. 6b). There are significant overlaps between these differentially expressed genes across different SLK cell types (Fig. 6c), indicating SLK isoforms are involved in similar network yet play distinct roles. Among them, L vs S and KO vs WT shared higher overlap than other comparisons (Fig. 6c). However, the genes differently expressed in L vs S is significantly enriched in pathways related to cancer development while the genes in KO vs WT may involved in other cellular functions (Fig. 6b). These findings further indicate the cancer-associate function of SLK attributes to the regulation in the splicing level other than the expression level. In addition, by comparing the SLK-L and SLK-S isoforms, we were able to account for the changes primarily caused by the splicing switch. Within these differential genes from SLK-L and SLK-S cell lines, we identified two EMT-related gene sets using GSEA. The GSEA also reveals slight enrichment of EMT gene sets in all comparisons (Fig. S5). Interestingly, SLK-S and SLK-KO exhibited a different trend of enrichment compared to the SLK-L. This observation suggests the involvement of different SLK isoforms in the process of cell migration.

**Figure 5.**
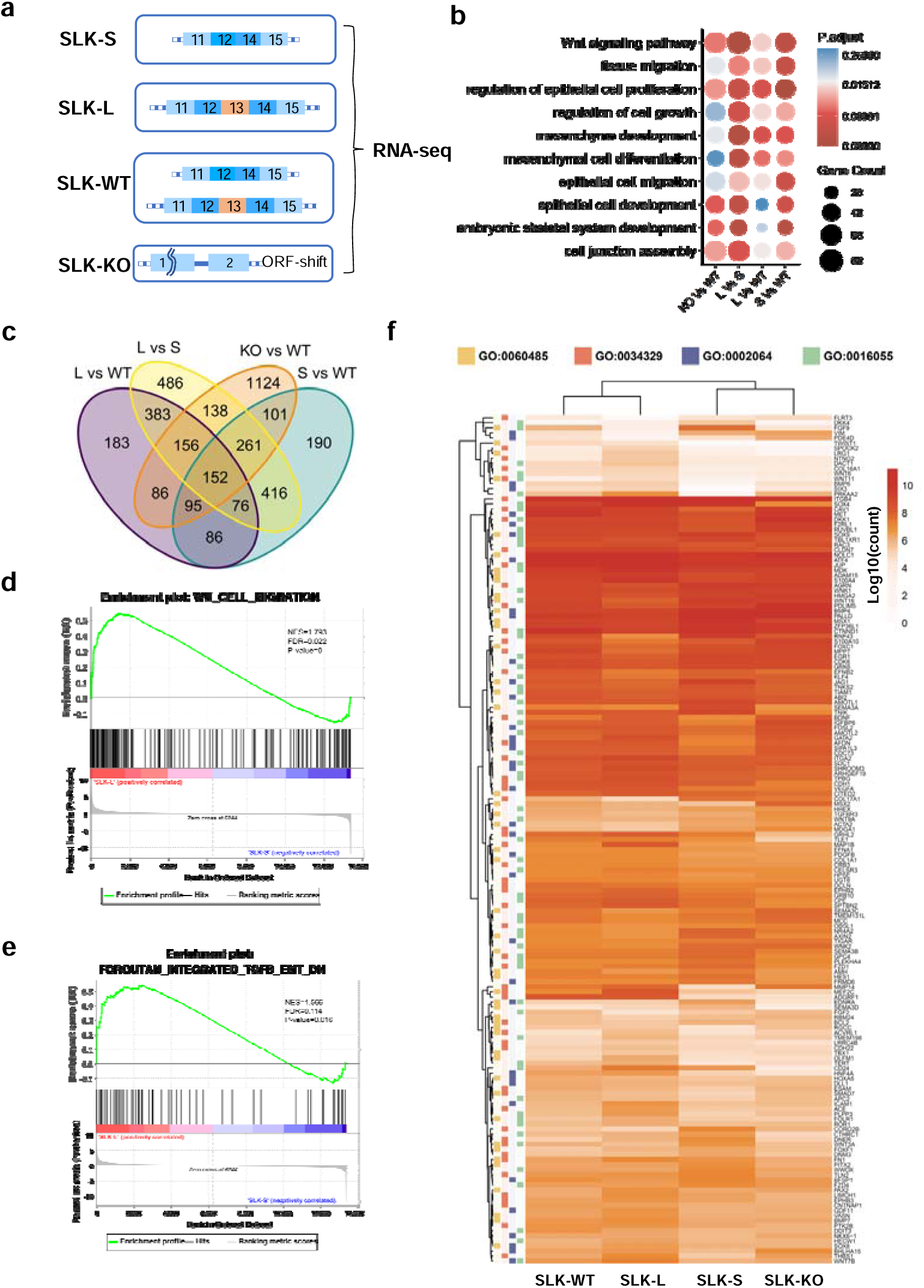
Rbfox2 regulates the splicing of SLK. **a,** Rbfox2 binding peaks was detected in front of the SLK exon 13^th^. **b,** The RIP assay showed the binding of Rbfox2 to SLK RNA. **c,** Rbfox2 expression level in tumors versus normal tissues across various cancer types from TCGA data. **d,** The minigene reporter of SLK wildtype and mutants were constructed. **e-f.** Rbfox2 regulates SLK splicing through its binding motif. **e,** Rbfox2 overexpression inhibited SLK minigene splicing. **f,** Rbfox2 knockdown promoted SLK minigene splicing. **g,** Rbfox2 regulates splicing of the endogenous *SLK* gene. RNA loading control: GAPDH; protein loading control: GAPDH or Tubulin. The chart represents mean±SD. *P<0.05, **P<0.01, ***P<0.001, ****P<0.0001. Two-way ANOVA was applied to analyze the difference among means in E and F while one-way ANOVA was used in G.

**Figure 6.**
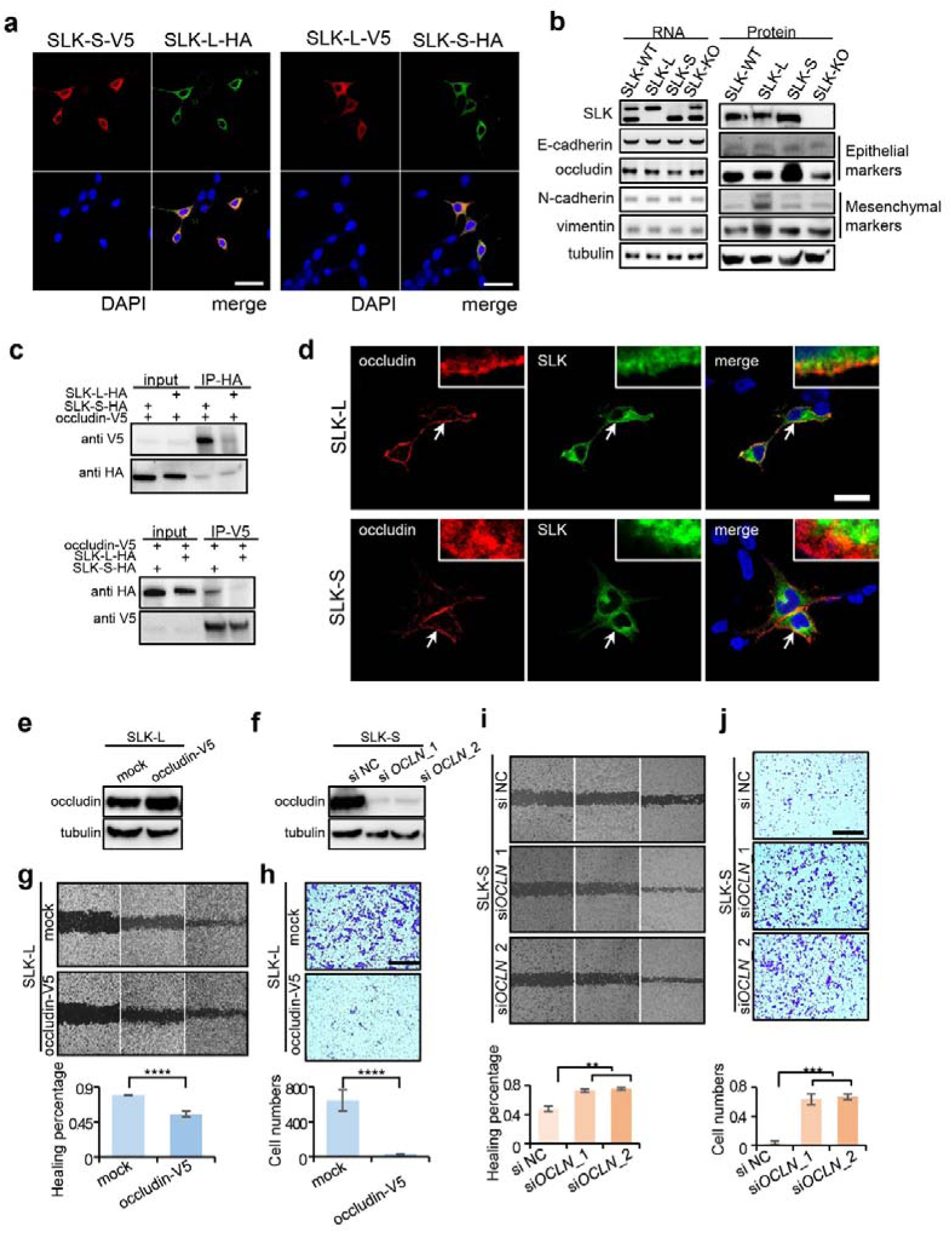
Switch of SLK isoforms change EMT-related genes. **a,** RNA-seq of indicated SLK cell lines. **b,** Gene ontology enrichment for differentially expressed genes from SLK-KO vs SLK-WT, SLK-L vs SLK-S, SLK-L vs SLK-WT and SLK-S vs SLK-WT. **c**, Overlap of differential genes from different comparisons. **d-e,** GSEA was performed to rank differentially expressed genes for SLK-L and SLK-S. **f,** Heatmap of overlapped differential genes in GO:0016055 “Wnt signaling pathway”, GO:0034329 “cell junction assembly”, GO:0002064 “epithelial cell development”, and GO:0060485 “mesenchyme development”.

Meanwhile, the Wnt signaling pathway can induce the expression of the key transcription factor Twist, which can repress the expression of epithelial markers such as E-cadherin, occludins, and claudins. On the contrary, it will induce expression of mesenchymal markers such as N-cadherin, fibronectin, and vimentin ^31^. Consistently, the expression of Twist cannot be detected in SLK-S cells but is moderately expressed in SLK-WT and SLK-L cells (Fig 6f), again demonstrated that the two isoforms play distinct roles in EMT and metastasis. To further identify potential gene targets of SLK in cancer metastasis, we retrieved all differentially expressed genes involved in cell junction assembly, Wnt signaling pathway, and regulation of epithelial cell proliferation, and found that many cell junction markers have been influenced. Collectively, RNA-seq of different SLK isoforms revealed the isoforms execute distinct functions in cellular pathways related to cancer in a systematic manner.

### SLK splicing switch influences cell migration in part through their different binding abilities to EMT and cell junction markers

Based on the experimental results and RNA-seq data, we consider that AS of SLK is closely intertwined with EMT and cell junction regulation, thus influencing cell migration. Considering that SLK-L has extra 31 amino acids which form a longer coiled-coil domain by the structural prediction and comparison by Alphafold (Fig. 1e), it indicates that the functional difference between the two isoforms attributes to the binding activity rather than the kinase activity. Therefore, there are two plausible explanations of how SLK isoforms can affect functions. First, the distinct isoforms may exhibit divergent cellular localization, enabling discrepant biological functions. Alternatively, the two SLK isoforms may have similar cellular localization, but act as antagonists and have differential interactions with corresponding targets, ultimately influencing distinct cascades. To investigate the mechanism, we first sought to determine whether the extra amino acids can affect the subcellular localization of SLK isoforms. By delivering different tagged SLK isoforms into the cells and detecting the immunofluorescence localization, we found both SLK isoforms locate near the cell membrane (Fig. 7a), suggesting their function difference is irrelevant to cellular localization.

**Figure 7.**
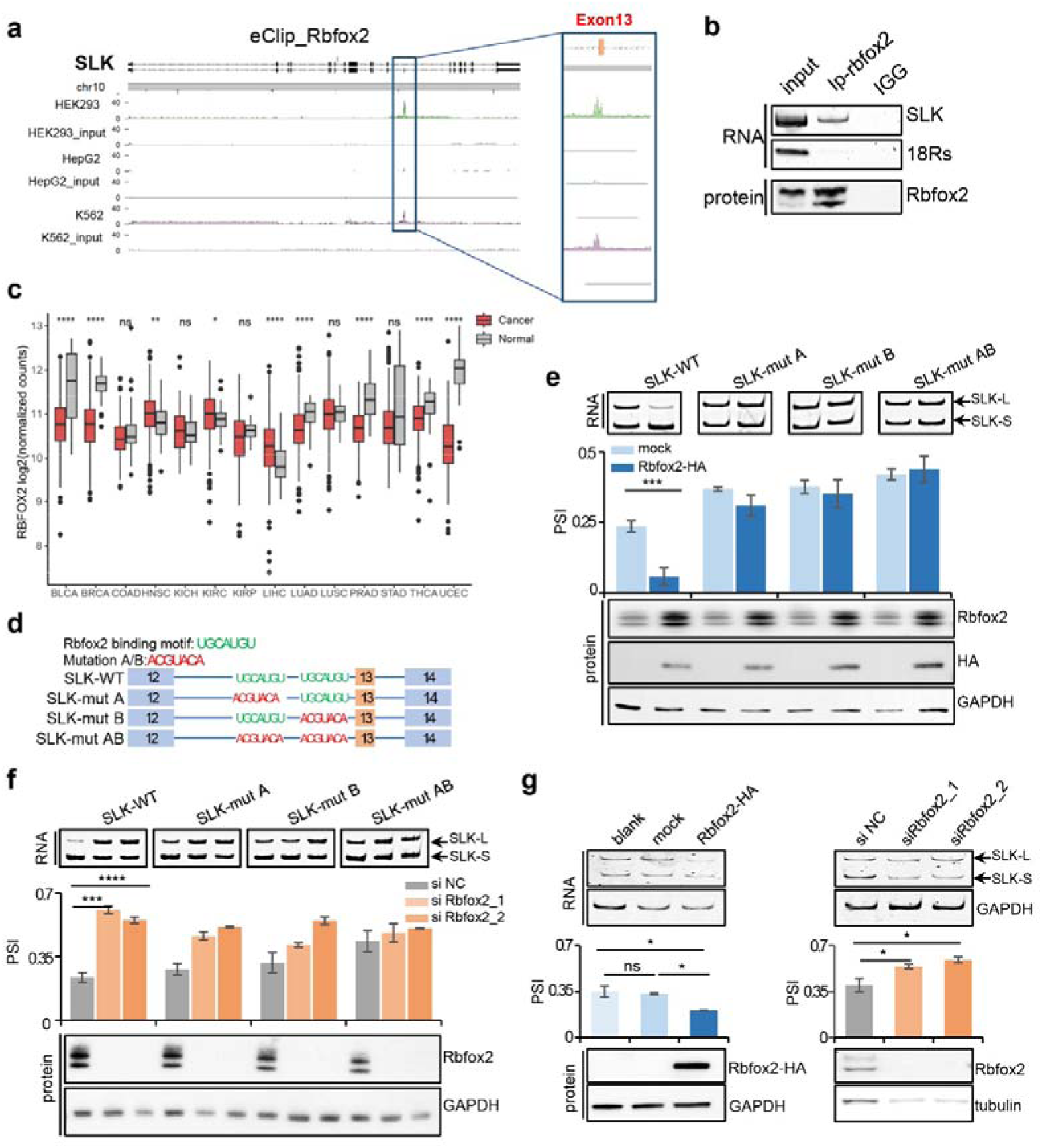
SLK isoforms can influence cell migration partially through their different binding abilities to cell junction markers. **a,** SLK isoforms demonstrate same cellular localization. **b,** The expression level of EMT markers in different SLK cell lines. **c,** SLK isoforms showed varied binding abilities with occludin. **d,** The subcellular localization of SLK and Occludin. Scale bar:20um. **e-j,** Regulation occludin affect cell migration and invasion in indicated SLK cell lines. **e,** Overexpression occludin in SLK-L cells. **f,** Knockdown the expression of occlidin in SLK-S cells. Cell migration (**g,i;** n=3), and invasion (**h,j;** n=3; scale bar:200µm).*P<0.05, **P<0.01, ***P<0.001, ****P<0.0001, ns:not significant. One-way ANOVA followed by Tukey’s test was used in **g-j**. The chart represents mean±SD.

As SLK isoform switch is closely linked to cell migration, we next tested several biomarkers of EMT such as E-cadherin, occludin, N-cadherin, and vimentin that were differentially expressed based on RNA-seq data to assess their expression in different SLK cell lines (Fig. 7b). As a result, occludin and vimentin were detected to be differentially expressed in SLK-L versus SLK-S stable cell lines. SLK-S cells have a higher expression level of occludin than SLK-L cells (Fig. 7c) whereas vimentin has a lower expression level in SLK-S cells than in SLK-L cells.

Occludin is considered a staple of tight junctions and performs vital functions in EMT, which contributes to cancer metastasis. To address whether SLK isoforms interact differentially with occludin, we test their interaction both through immunoprecipitation and localization. Immunoprecipitation showed that SLK isoforms had different binding capabilities with occludin. The SLK-S had a stronger binding ability to occludin, while the SLK-L was barely bound to occludin (Fig. 7c). Furthermore, the immune colocalization demonstrated that both SLK isoforms and occludin localized around the cell membrane (Fig. 7d), indicating the possible interaction between SLK and occludin.

Given cancer cells with low occludin expression are more likely to migrate and invade ^32^, we assess the relationship between occludin expression level and cell development in this study by overexpressing occludin in SLK-L cells (Fig. 7e) or knockdown occludin in SLK-S cells (Fig. 7f). Consistently, the overexpression of occludin in SLK-L cells could inhibit cell migration and invasion (Fig. 7g), whereas knockdown occludin in SLK-S cells promoted cell migration and invasion (Fig. 7j). The findings suggest that SLK-L could function as an oncogenic protein by disrupting the interaction between SLK-S and occludin. The overexpression of the SLK long isoform may destabilize tight junctions (TJs), resulting in irregular cell fate. These results support the notion that high levels of SLK-L expression are associated with carcinogenicity and the promotion of cancer metastasis, providing valuable insight into this phenomenon.

### The RNA binding protein Rbfox2 regulates SLK alternative splicing

We next proceeded to investigate the regulatory mechanism responsible for the AS of SLK. In addition to the regulation by the core spliceosome ^33^, AS is also controlled by the cooperation of multiple regulatory *cis*-elements that recruit *trans*-acting splicing factors to affect proximal splice sites (SS) through multifarious mechanisms ^33–36^. Since the AS of SLK exhibited significant effects on cancer development, we aim to investigate how SLK splicing is regulated. We downloaded RBP binding peaks from the ENCODE eCLIP-seq dataset ^37^ and identified the potential splicing factors that could regulate the splicing of SLK 13^th^ exon. As a result, three splicing factors were discovered to have binding peaks near the 12^th^ intron and the 13^th^ exon of SLK, including Rbfox2, QKI, and HNRNPA1 (Fig. 5a and S4a). We overexpressed these splicing factors to test their function. Rbfox2 overexpression is found to change the AS of SLK while QKI and HNRNPA1 exhibited less influence (Fig. S4b). In addition, the RNA immunoprecipitation (RIP) assay confirmed the binding of Rbfox2 to SLK RNA (Fig. 5b). Rbfox2 is a member of the RNA-binding Fox proteins (Rbfox) family which regulates splicing by binding to the motif (5’-UGCAUGU-3’) on the pre-RNAs ^38–41^. The expression of Rbfox2 is found to decrease in the cancer types in which the PSI of SLK increased (Fig. 5c).

To test whether Rbfox2 inhibits SLK splicing by binding to its pre-RNA, we checked the sequence of SLK RNA and found two Rbfox2 binding motifs in the 12^th^ intron. Then we constructed SLK splicing reporters with the wildtype or mutated Rbfox2 binding motifs to assess the regulatory mechanism of Rbfox2 (Fig. 5d). Subsequently, we tested the splicing of SLK reporters in cells undergoing Rbfox2 overexpression or knockdown. As expected, in Rbfox2-overexpressed cells, AS of the wild-type SLK reporter tended to skip the 13^th^ exon, producing more SLK-S. However, the skipping ability of the mutated reporter was attenuated, leaving the splicing ratio similar to the mock control (Fig. 5e). On the contrary, the SLK-WT reporter tended to retain the 13^th^ exon in Rbfox2 knockdown cells, resulting in the AS shift towards SLK-L whereas the splicing of mutated reporters remained unchanged with Rbfox2 deficiency (Fig. 5f). The reporter gene assay confirmed that Rbfox2 regulates SLK alternative splicing by binding to the motif sequence 5’-UGCAUG-3’ in the 12^th^ intron to promote the skipping of SLK 13^th^ exon, resulting in lower PSI and decreased SLK long isoform.

In addition, we tested the regulation of Rbfox2 to endogenous AS of SLK. Rbfox2 was either overexpressed or knocked down in cancer cells to regulate the AS. The results were consistent with that of the SLK reporter. Endogenous 13^th^ exon of SLK skipped after overexpression of Rbfox2 as compared to nonspecific and negative controls (Fig. 5g, left), whereas it was included after knockdown of Rbfox2 by siRNA (Fig. 5g, right). Collectively, the results demonstrated that Rbfox2 can regulate SLK AS by binding to its recognition motif on the SLK 12^th^ intron.

## Discussion

More than 90% of human genes undergo alternative splicing ^42^. After eliminating noises caused by tissue-specific splicing, numerous gene-splicing alterations have been detected in a large amount of cancer patients ^9^. Abnormal splicing is considered a molecular hallmark of cancer and thus worth efforts in investigating the underlying mechanisms, the cellular consequences, and the possibility of manipulation, for cancer treatment. In this study, we have demonstrated that the significant and extensive alterations across multiple cancer types are involved in actin filament organization, cell junction assembly and cell-substrate adhesion, which suggests a universal functional and mechanistic implications. We demonstrated SLK AS is one of the crucial splicing events that are dysregulated in cancer and splicing switch to SLK-S results in cancer inhibition.

As a protein kinase, previous studies of SLK focused on its catalytic activities. Recently, a few studies have demonstrated that some cellular functions of SLK appears to be kinase activity independent, suggesting that it may be an important scaffold for signal transduction pathways ^14^. In some cases, overexpression of scaffolding proteins often leads to the titration of signaling components, while their knockdown interferes with the formation of signaling complexes, resulting in similar phenotypes ^43^. Our study of the AS of SLK by coincidence shed light on these discoveries, by implying the possibility that this may not be a regulation of the splicing level rather than the expression level.

Interestingly, in the animal experiments of testing AS of SLK, our study found that SLK knockout cell lines showed similar proliferation rate in the nude mice compared to SLK-WT though knocking out SLK is embryonic lethal. On one hand, it indicates an undiscovered complementary mechanism upon SLK knockout. On the other hand, it implies that targeting the expression of SLK may be therapeutically ineffective. In contrast, we discovered the SLK-S cells produced smaller subcutaneous tumors in tumorigenesis experiments and significant less metastasis burden than other SLK groups. Collectively, these findings indicate that AS of the *SLK* gene has the potential to be a target for tumor diagnosis and therapy, which need further validation.

We have discovered that occludin may have different binding abilities to different SLK isoforms, However, the precise mechanism of how they interact with each other remained for further exploration. Most occludin functions were initially attributed to conformational changes following selective phosphorylation ^44^, hence more solid validations between SLK AS and occludin are required.

The mechanisms of the SLK splicing were found to be comprehensive. The AS of SLK was proved to be regulated by Rbfox2 through the binding motif 5’-UGCAUG-3’ on SLK pre-RNA. However, the expression of Rbfox2 was not simultaneously altered in some cancer types compared with SLK splicing (Fig. 5c), implying the AS of SLK is regulated in a complicated manner. Moreover, though overexpression of Rbfox2 can suppress SLK exon 13^th^ splicing, its regulation did not exhibit significant tumor inhibition in vitro (data not shown). In addition, Rbfox2 is a ubiquitous RNA binding protein that have various RNA targets other than SLK, thus therapeutics strategies targeting Rbfox2 to regulate SLK alternative does not have promising outlook. Nevertheless, Rbfox2 form a large assembly of splicing regulators (LASR) ^45, 46^, thus the components of LASR also have the potential in affecting SLK splicing. In summary, the expression levels of SLK isoforms showed drastic variations in tumor cells and normal cells. We found that different isoforms of the *SLK* gene had different biological functions. SLK long isoform is crucial for cancer maintenance and development. SLK isoforms might affect cell functions by the different binding abilities of occludin. Rbfox2 inhibited SLK exon 13 splicing through its binding motif. Collectively, our study uncovers the biological consequences and underlying mechanisms for one of the most mis-spliced genes in cancer, highlighting its potential significance in cancer diagnosis and treatment. By taking *SLK* as an example, we hope to draw attention to the importance of splicing regulation in cancer.

## Materials and Methods

### Plasmid construction

The CDS sequences of SLK-L and SLK-S were amplified from the cDNA of HCT116 cells. The two SLK isoforms were cloned into the pcDNA3.1 overexpression vector with the HA or V5 tag sequence. The CDS regions of SLK isoforms with the HA or Flag tag or the firefly luciferase CDS were inserted after the CMV promoter of the pCDH backbone plasmid to create lentivirus vectors, respectively. The Rbfox2 CDS region with the HA tag was cloned into the pcDNA3.1 backbone to produce Rbfox2 overexpression vectors.

For the SLK minigene reporter, we cloned SLK exon 12, 13, 14, a 337-nt sequence with 5’-SS, a 290-nt sequence containing branch point and 3’-SS of intron 12, and a fragment of intron 13, into the pcDNA3.1 plasmid backbone. The SLK mutation vectors were constructed by replacing the Rbfox2 binding motif 5‘-TGCATGT-3’ with 5‘-ACGTACA-3’.

To generate stable cell lines with SLK-S or SLK knockout, we designed guide RNAs (gRNAs) targeting specific sites of SLK genome DNA and cloned them into the CRISPR-Cas9 plasmid. For cancer cell lines selectively expressed SLK-L, an additional template of exon13-14 or 12,14 were supplied as the donor DNA at the target site to allow the cells to repair the double-strand break through homology-directed repair (HDR) pathways (Fig. 2a, Fig. 3a and Table S1).

### Co-immunoprecipitation

293T or HCT116 cells were seeded in a 15-cm dish and transfected with HA-tagged SLK isoforms and V5-tagged candidate proteins. The PBS-washed cells were used after transfection for 48 h. The lysis buffer (P0013, Beyotime) was then added into the cells and the supernatants were collected after centrifuging at 15000 rpm at 4 °C for 30 min. Next, 10% supernatant was retained as input and the pallet was resuspended. Primary antibodies (HA-Tag Mouse mAb #2367 and V5-Tag Rabbit mAb #13202, CST) were added and incubated overnight. The antibodies were diluted at the ratio of 1:100. Subsequently, 50 µl protein A/G agarose beads were added to each tube and incubated at room temperature for 1 h. Following centrifugation at 2500 rpm for 5 min, the supernatants were removed, and the agarose beads were resuspended in 500 µl IP buffer and washed at room temperature 3 times. After the washing step, 50 µl of 2×SDS protein loading buffer was added, and the denatured proteins were obtained by incubation at 98 °C for 10 min.

### RNA Immunoprecipitation

Cells grown in a 15-cm dish (90% confluence) were digested with 3 mL of 0.25% trypsin at 37 °C for 2 min. Next, 5 mL of complete medium was added to the dish to terminate digestion. The collected cells were subjected to RNA immunoprecipitation following the Abcam protocol (https://www.abcam.com/epigenetics/rna-immunoprecipitation-rip-protocol).

### Cancer cell development and function

Cell proliferations were analyzed by cell counting or CCK-8 assay. For cell counting, 5000 cells were seeded per well on 24-well plates, and the cell number was counted at 24, 48, 72, and 96 h after incubation. For the CCK-8 assay, 1000 cells were seeded per well on 96-well plates, and the Cell Counting Kit-8 (CCK-8, 40203ES60, Yeason) was used to determine the cell proliferation.

A colony formation assay was carried out to test the tumor formation ability of cancer cells. In brief, 2000 cells were seeded onto a 6-well plate in a complete medium with 10% FBS for one week. The cells were then dyed with crystal violet for 30 min before washing with PBS. The plating efficiency was calculated using colony numbers divided by seeded cell numbers.

Cell migration ability was measured through wound healing experiments. Cells were seeded onto a 6-well plate. After reaching 90% confluence, the cells were scratched with a 200-µl pipette tip. The images were captured at 0, 12, 24, and 48hr after scratching. The average value of wound closure was quantified by Image J software. Experiments were performed in triplicates.

Transwell invasion assay was used to determine the invasion ability of cells. Matrigel Matrix (356234, Corning) was thawed at 4 °C and mixed with medium without FBS at the ratio of 1:8. Next, 50 µl of the mixture was added to each Transwell and incubated at 37 °C for 1 h. Subsequently, 1×10^5^ cells (2×10^5^ cells in Fig 3F) were seeded in each Transwell in 200 µl medium without FBS, and 500 µl medium with 10% FBS was added as an inducer in the 24-well-plate. The cells were cultured at 37 °C for 24 h. The matrix and media were removed, and the cells were incubated with 4% paraformaldehyde (PFA) for 30 min after washing with PBS. After removing PFA, the cells were washed with PBS twice. The cells were then stained with crystal violet for 30 min and de-stained with PBS twice before imaging. Cells were counted from the captured images by using Image J software. Each experiment was repeated three times.

### Lentivirus packaging and infection

HEK 293T cells were plated at 50% confluence in a 10-cm dish before transfection with plasmids. The envelope plasmid pMD2.G, the packaging plasmid psPAX2, and the transfer plasmids were co-transfected into HEK 293T cells by Lipofectamine 3000. The mediums with lentivirus from transfected cells were collected after 48 h of transfection and filtered through a 0.22-µm membrane.

### Immunofluorescence assay

293T cells were seeded in a Nunc^TM^ dish (150680, Thermo) and transfected with HA-tagged SLK isoforms and V5-tagged occludin (for Fig 6C) or SLK different isoforms with HA or V5 tag (for Fig S5). Cells were incubated with 4% PFA for 1h. After being washed with PBS, cells were incubated with 1mL 0.25% Triton^TM^ X-100 (9036-19-5, Meck) per dish for 10 min. Then cells were washed with PBS twice and incubated with HA and V5 antibodies (dilution ratio: 1:1000) at 4°C overnight. Alexa Fluor® 594 (ab 150080, Abcam) and Alexa Fluor® 488 (ab 150077, Abcam) were Incubated with cells at room temperature for 1h. The nuclei were stained with DIPA (P36985, Invitrogen™) and observed and photographed under Zeiss LSM880.

### Xenograft tumor formation

We conducted two biological replicants of tumorigenesis experiments in nude mice. In the first replicant, 5×10^6^ cells were resuspended in PBS and injected subcutaneously into a BALB/c-nude mouse (10 mouse/group). The body weight and tumor volume were measured twice a week after injection. In SLK-WT and SLK-L groups, several mice were sacrificed before the endpoint due to the fast-growing tumors and the following tissue rupture. In the second biological replicant, 1×10^6^ cells were resuspended in PBS and injected subcutaneously into a BALB/c nude mouse. The tumor volume was measured every three days. The experiment was terminated 33 days after injection. The tumor volume was calculated as volume (mm^3^) = L ×W^2^/2, where L is the largest diameter and W is the smallest diameter of the tumor. After the experiment, the mice were euthanized, and the tumor was dissected.

### Tail vein metastasis experiment

Tail vein metastasis experiments were carried out in both BALB/c nude mice and C-NKG mice at Cyagen Co., Ltd (SuZhou, China). Briefly, luciferase-labeled HCT116 cells (SLK-WT, SLK-L, or SLK-S) were resuspended in 100 μL PBS at concentrations of 1.0×10^6^ or 2.0×10^6^ and injected into the lateral tail veins of seven-week-old C-NKG or BALB/c nude female mice between 18-22 g, respectively. The mice were anesthetized and injected intraperitoneally with D-luciferin (Yeasen, China) for a dose of 150 mg/kg on the indicated day post-injection. Metastasis was quantified by total flux (photons/second) of bioluminescence in the lungs on a PerkinElmer IVIS Lumina III *in vivo* imaging system.

### Bioinformatic analyses

To identify the cancer-related alternative splicing from TCGA, we downloaded the analyzed data from the published database (IDEAS), which calculate the PSI value with rMATs. We filtered out cancer types with fewer than 20 normal tissue counterparts to ensure robust and reliable estimation of PSI changes. The Wilcox test was applied to determine the significance between cancer and normal groups. For alternative splicing events shown in Figure S1a, we only included the significantly changed splicing events with ≥10 read counts, and ≥5 observations in normal samples. We downloaded the level 3 RNA-seq data from TCGA (http://gdac.broadinstitute.org/), and the splicing events shared by more than 10 cancer types were analyzed in the GO enrichment network.

### CLIP-seq analysis

The processed signal files for eCLIP-seq of all RBPs were downloaded from the ENCODE consortium. To visualize binding peaks of the indicated RBPs, we utilized the R packages ‘Gviz’ and ‘rtracklayer’. For Figure 5a, the HepG2 and K562 data were downloaded from the ENCODE consortium, and the HEK293 data were obtained from the GEO database (GSE77629). All eCLIP-seq data were aligned to GENCODE hg38 v39 using STAR and then visualized using karyoploteR.

### RNA-seq analysis of SLK cell types

For the RNA-seq analysis, we used trim galore to remove the adapter from sequencing data, followed by gene expression calculation using RSEM based on GENCODE hg38 v39. Gene count data were imported using the tximport package and normalized by the TMM method from NOISeq. NOISeq was used to obtain differentially expressed genes from different cell lines, with q=0.8 and |M| > 0.5 using in NOISeq analysis. ClusterProfiler package was used to perform GO enrichment analysis, and ggplot2 was applied to visualize the results. The Venn Diagram was constructed using the ‘VennDiagram’ package. Gene sets that were differentially expressed in both L vs S and L vs WT, L vs S and S vs WT, and L vs S and KO vs WT were further investigated as potential targets. The gene expression profiles of these genes were plotted using ‘heatmap’.

### Structure prediction of SLK-L and SLK-S

Parafold ^47^, a paralyzed accelerated version of alphafold2 ^24^ was applied to predict the structure of SLK-L and SLK-S. The input is the RNA sequence of the open reading frame of each isoform. Deepmind monomer model sets were employed for generating the prediction. The top 5 ranked results were investigated according to Alphafold default evaluation.

### Data Availability

eCLIP data of RBPs and the processed signal files (in bigwig format) were downloaded from ENCODE consortium (https://www.encodeproject.org). RNA-seq data during the current study are available in the NODE data depository. Custom computer code and original data are available upon request.

## Acknowledgments

We thank Prof. Yong-Bo Wang of Fudan University and Prof. Yang Wang of Dalian Medical University for the manuscript revision. We thank Ms. Yun Jiang, Dr. Yujie Chen, Ms. Nianhao Cheng, and Ms. Shuang Han for their assistance with the experiments. We are grateful for the experimental support of the UliSchwarz Quantitative Biology Core Facility in Bio-Med Big Data Center, SINH, CAS. This work is supported by the National Natural Science Foundation of China (32171294) and Shanghai Municipal Natural Science Foundation (20ZR1467300) to H-H. Wei. It is also supported by the National Natural Science Foundation of China (31730110, 91940303, and 32030064), the National Key Research and Development Program of China (2018YFA0107602), and the Strategic Priority Research Program of the Chinese Academy of Sciences (XDB38040100) to Z. Wang.

## Author contributions

H-H. Wei and Z. Wang conceived the project. H-H Wei designed the experiments and interpreted the results. Y-Q. Yang, H-H. Wei and J. Sun carried out molecular, biochemical, and cell function experiments. Y. Hu and S-R. Zhang analyzed the eCLIP-seq, TCGA data, and RNA-seq data and performed other bioinformatic analysis. H-H. Wei was responsible for the tail vein injection experiment. W-J. Zhang and Y. Sun performed the mice xenograft experiment. J-F. Li and J-W. Guan performed protein structure prediction. X-Y. Feng generated the CRISPR HCT116 cell lines. H-H. Wei, Y-Q. yang and Y. Hu wrote the manuscript. Y. Yang helped to interpret the data.

