## Supplementary Figures for "Extensive Dysregulation of SLK Splicing in Cancers Impacts Metastasis"

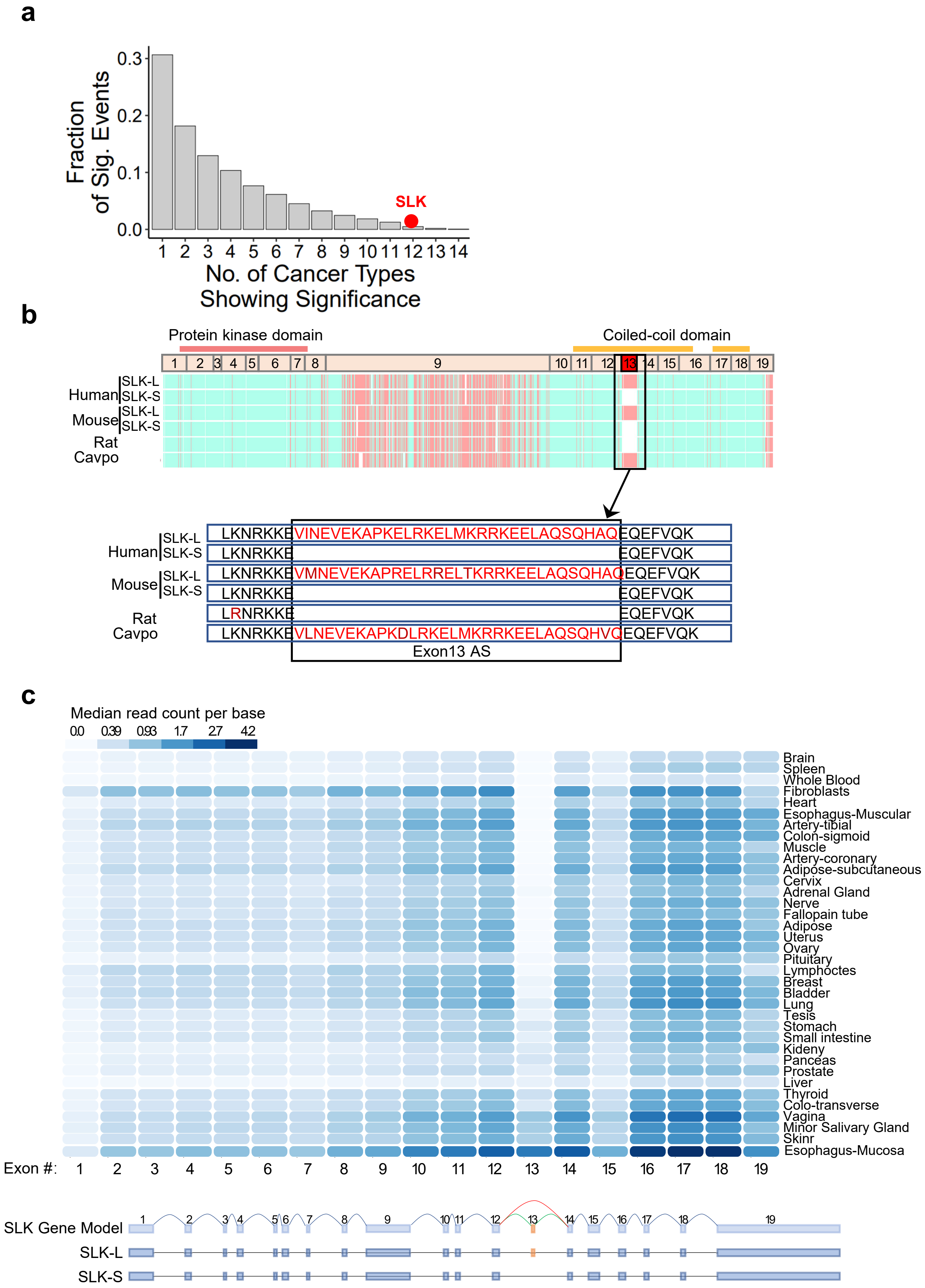


**Figure S1 (related to Fig. 1) a,** Fractions of alternative spliced genes in different cancer types. **b,** The SLK splicing is conserved across mammals. **c,** The expression of SLK isoforms in multiple tissues.

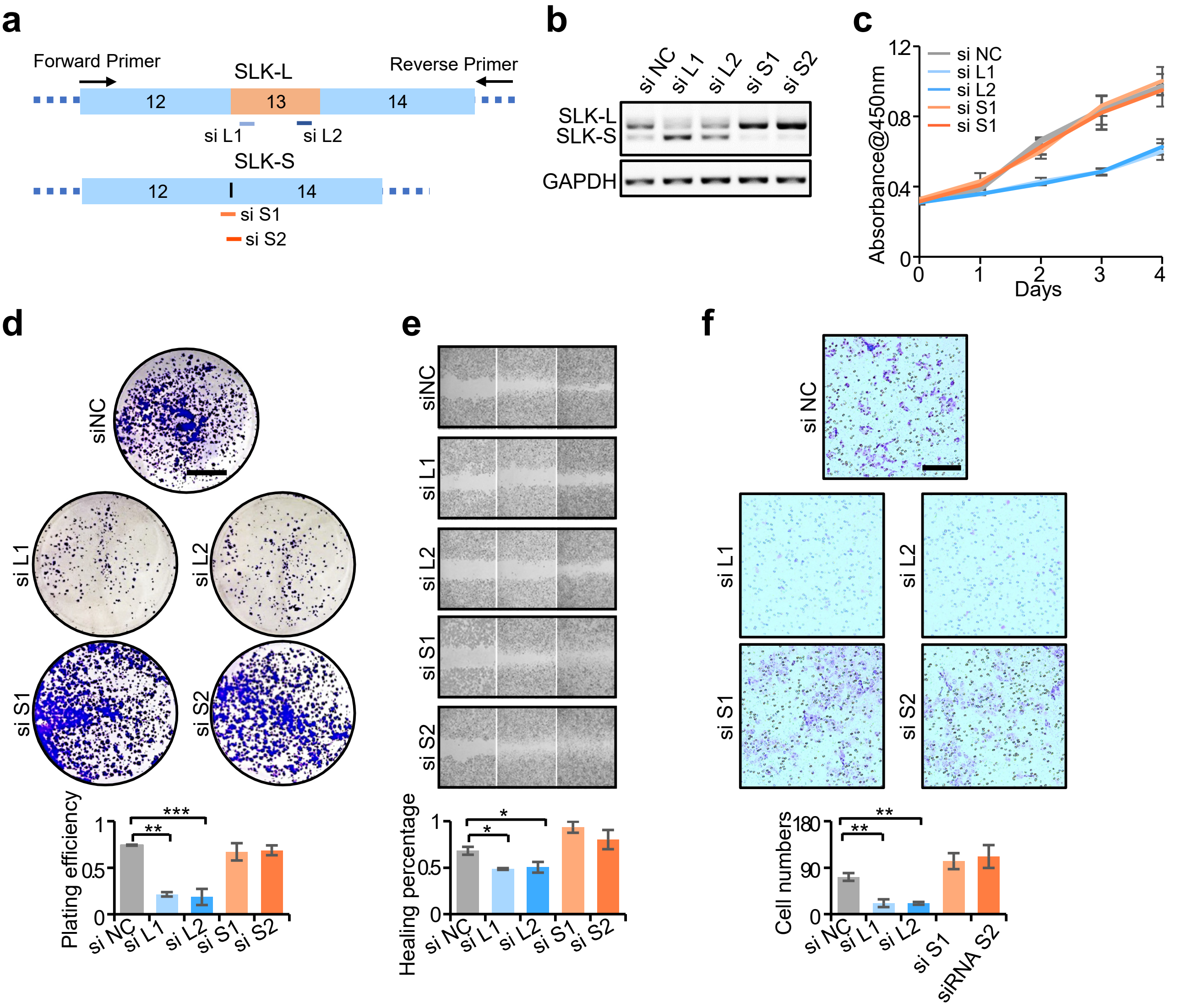


**Figure S2 (related to Fig. 2)** siRNA treatment affected cancer progression by altering the expression levels of SLK isoforms (related to Fig.2). **a,** The design of siRNA targeting SLK-L or SLK-S. **b**, Knockout efficiency of the siRNAs was determined by RT-PCR. **c-f,** Cell functions were tested according to Fig.2. The mean ± SD were plotted, with p values calculated by Student's *t* test.

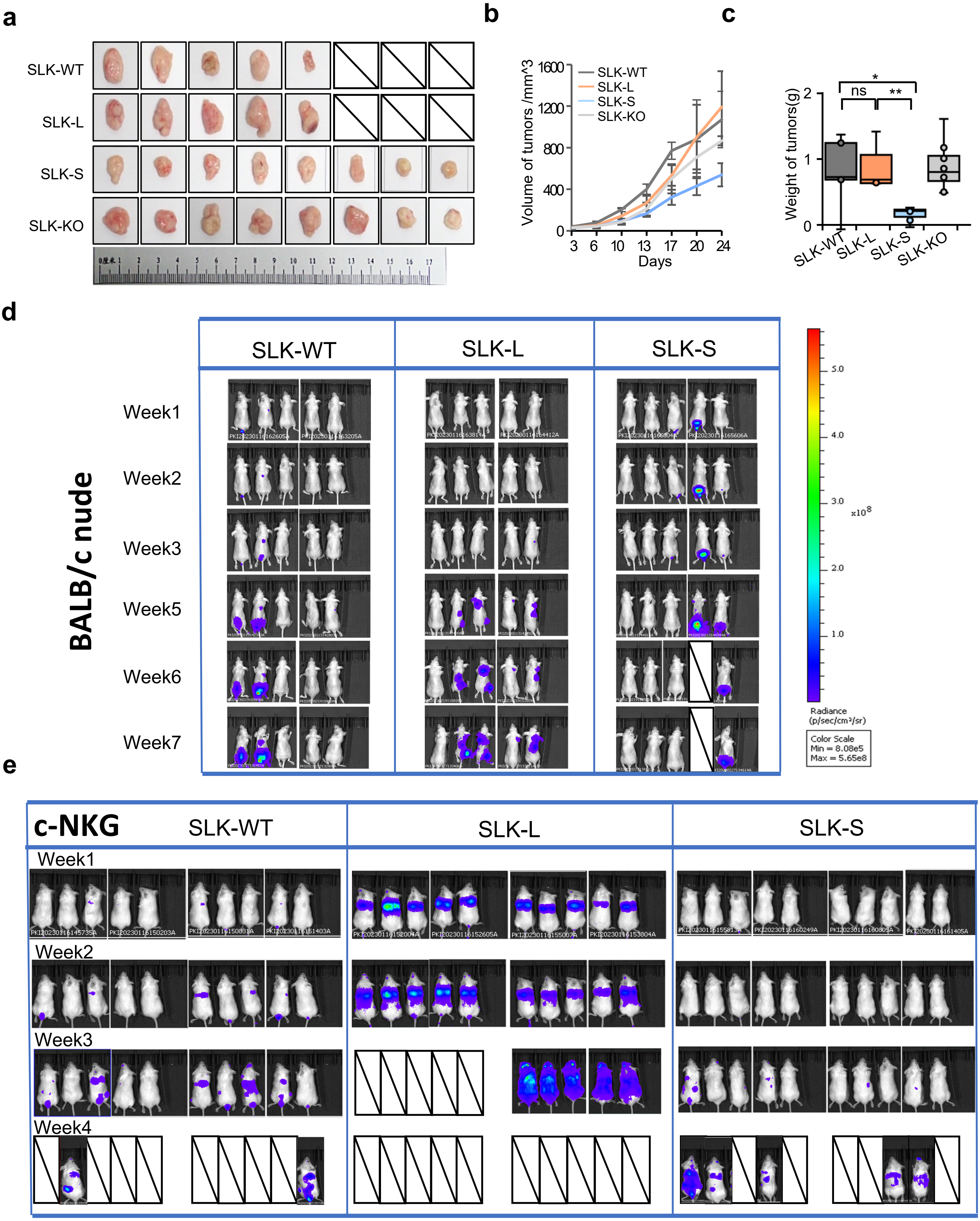


**Figure S3 (related to Fig. 4)** Different groups of SLK cells exhibited distinct impacts on cancer development (related to Fig. 4). **a-c,** Xenograft tumors were generated in BALBc/nude mice by injection with SLK-WT, SLK-L, SLK-S, or SLK-KO cells ($5 \times 10^6$ cells/mouse), respectively. Tumor volume and weight were quantified and demonstrated in **b** and **c**, with *p* values calculated by Dunnett's test. **d-e,** Metastasis of SLK cell groups was measured in two mouse strains. Different SLK cells were injected in the lateral tail vein of C-NKG mice (d) and BALB/c nude mice (e), and the bioluminescence was detected in the indicated time post injection. Diagonal grid represented the death of experimental mice.

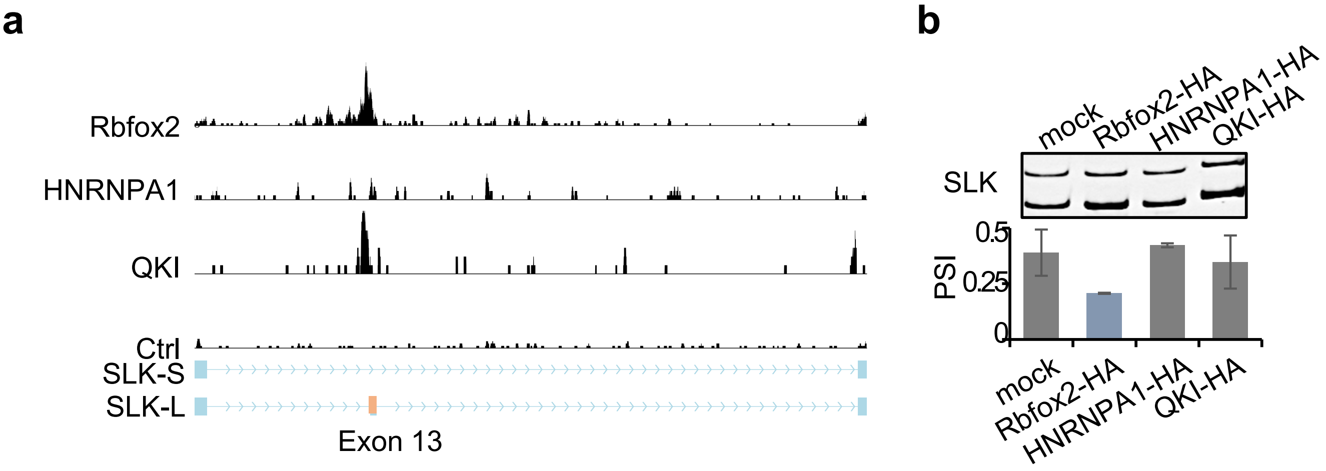


**Figure S4 (related to Fig. 5)** The candidate SLK splicing factors were discovered and validated. **a,** Various splicing factors exhibited binding peaks near the SLK 13 exon. **b,** The splicing of SLK was demonstrated by semi-quantified RT-PCR upon treatment of selected splicing factors.

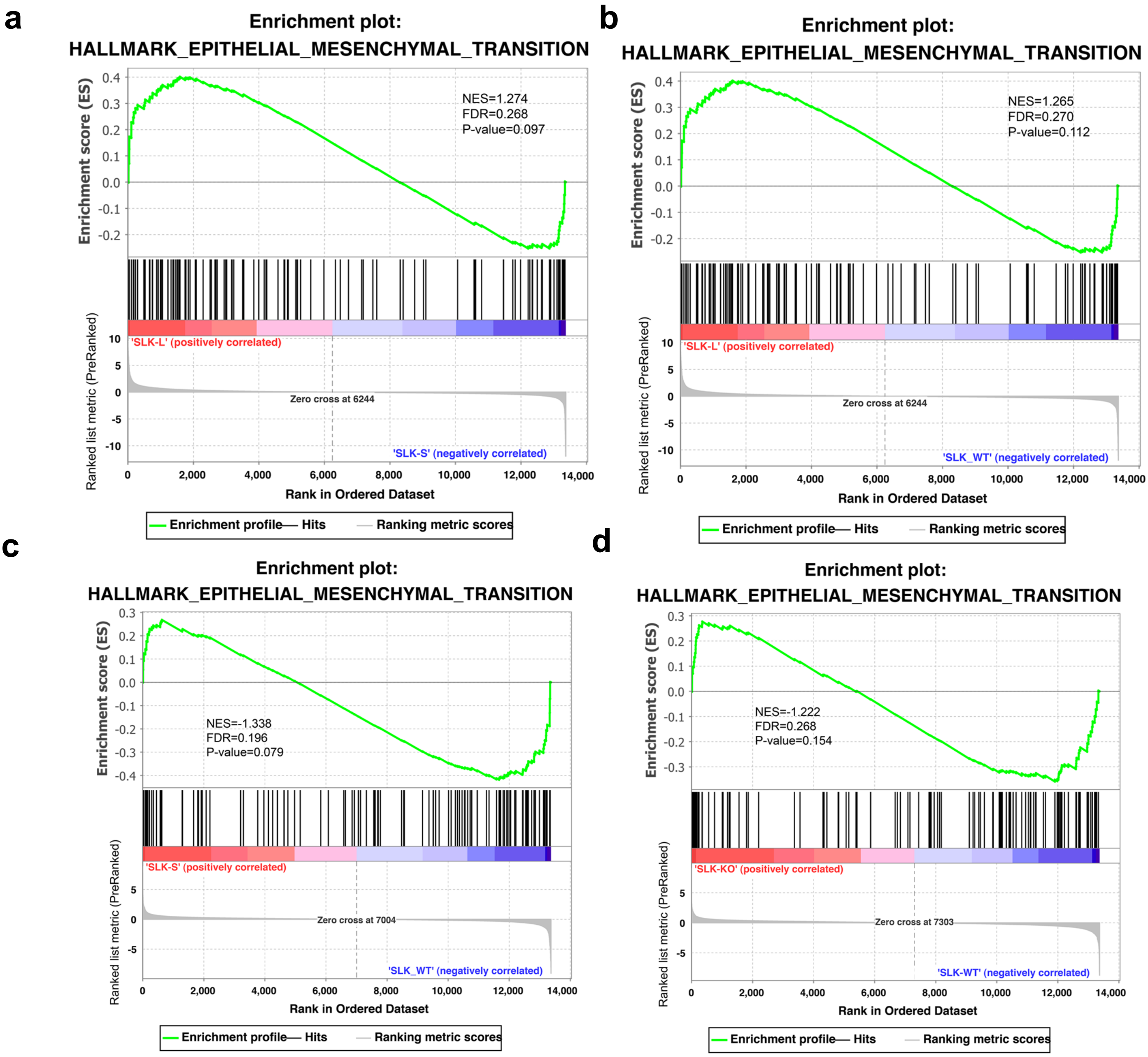


**Figure S5 (related to Fig. 6) a-d,** GSEA for differential genes in hallmarks of EMT. **a,** SLK-L vs SLK-S **b,** SLK-L vs SLK-WT **c,** SLK-S vs SLK-WT **d,** SLK-KO vs SLK-WT
